# Protein domain embeddings for fast and accurate similarity search

**DOI:** 10.1101/2023.11.27.567555

**Authors:** Benjamin Giovanni Iovino, Haixu Tang, Yuzhen Ye

**Affiliations:** Luddy School of Informatics, Computing and Engineering, Indiana University, 700 N. Woodlawn Avenue, Bloomington, IN 47408

**Keywords:** protein language model (PLM), ESM-2, domain segmentation, recursive cut, discrete cosine transformation (DCT), DCT fingerprint, homology detection

## Abstract

Recently developed protein language models have enabled a variety of applications with the protein contextual embeddings they produce. Per-protein representations (each protein is represented as a vector of fixed dimension) can be derived via averaging the embeddings of individual residues, or applying matrix transformation techniques such as the discrete cosine transformation to matrices of residue embeddings. Such protein-level embeddings have been applied to enable fast searches of similar proteins, however limitations have been found; for example, PROST is good at detecting global homologs but not local homologs, and knnProtT5 excels for proteins of single domains but not multi-domain proteins. Here we propose a novel approach that first segments proteins into domains (or subdomains) and then applies the discrete cosine transformation to the vectorized embeddings of residues in each domain to infer domain-level contextual vectors. Our approach, called DCTdomain, utilizes predicted contact maps from ESM-2 for domain segmentation, which is formulated as a *domain segmentation* problem and can be solved using a *recursive cut* algorithm (RecCut in short) in quadratic time to the protein length; for comparison, an existing approach for domain segmentation uses a cubic-time algorithm. We showed such domain-level contextual vectors (termed as *DCT fingerprints*) enable fast and accurate detection of similarity between proteins that share global similarities but with undefined extended regions between shared domains, and those that only share local similarities.

## Introduction

Homology detection is one of the fundamental computations in biology due to it’s role in helping determine protein function and structure. Every homology detection task begins with a protein sequence of interest that is queried against a collection of sequences with the goal of returning the most similar sequence as the top result. Despite the simplicity of this task in its conception, it can be incredibly difficult in practice due to the lack of similarity between two proteins that are considered homologous. Sequences with less than 10-12% similarity (in terms of character identity) have been found to contain similar structures [1], and thus it is crucial to detect proteins, or “remote homologs”, within this realm. Many methods have been developed to accurately and efficiently perform this task, ranging from simple (albeit heuristic heavy) sequence-sequence comparisons such as BLAST [2], to profile methods that consider groups of proteins like PSI-BLAST [3] and CS-BLAST [4], to profile hidden Markov models (pHMMs) based on probabilistic models like HMMER [5] and HHsearch [6] which are considered state-of-the-art for homology detection.

Recent methods, such as knnProtT5 [7] and PROST [8], have been developed using contextualized embeddings generated by neural networks, such as neural protein language models (pLMs), for homology detection. pLMs are trained for the purpose of learning about the nature of proteins beyond their sequence representation [9]. Because the sequence of a protein is constrained by the structure it folds into, each residue in a protein has more meaning than its character identity as each residue plays a role in the protein’s overall structure and function. This contextual information is learned by pLMs when they are trained on large sequence databases, like UniProt [10] and BFD [11] [12]. Many different pLMs have been trained and they can take on various architectures. ProtTrans [9], which contains a host of different models, was trained with the goal of producing informative embeddings as input for downstream tasks, such as predicting secondary structure and sub-cellular localization. ESM-2 [13] was trained with the purpose of producing embeddings that facilitate protein structure prediction based off sequence alone. Embeddings from both models have been successfully applied to many tasks, including homology detection, such as knnProtT5 [7] using embeddings from ProtT5 to perform nearest neighbor searches and PROST [8] using embeddings from ESM-1b to perform similarity searches.

One tradeoff with using pLM embeddings to represent protein sequences is the increase in dimensionality compared to the original character representation. For example, by embedding each position in a protein sequence N residues long, each position will be replaced by a vector of M length, where M can be in the thousands depending on the pLM used, so the protein sequence is now represented as a *N*×*M* matrix. Even when using simple distance metrics to calculate the difference between the residual embeddings, it will be computationally demanding to compare a query protein represented as such against every similarly represented protein in a database, because proteins are of various lengths and residual embeddings need to be aligned. Protein-level embeddings can offset this increase. A typical approach of deriving a protein-level embedding for a protein is to use the mean of the embeddings of all its residues. The mean embeddings have been used in applications, such as in knnProtT5 for remote homology detection using nearest neighbor search on protein-level embedding spaces [7]. The advantage of using protein-level embeddings is that proteins are now represented as a single vector of a fixed length, so similar proteins can be found by searching for proteins sharing similar embeddings, which can be computed quickly (L1-distance or other metrics can be used). When combined with other methods (MMSeq2 [14] or Smith-Waterman [15]), knnProtT5 achieved comparable performance with sequence-based approaches for homology detection, but it was not competitive for comparison of multi-domain proteins [7].

Another technique that has been applied with success is the iDCT vector quantization of embeddings. Introduced by WARP [16] and used in PROST [8], this method uses the Discrete Cosine Transform (DCT) [17], a commonly used technique in image and video compression, to reduce protein embeddings to smaller dimensions while maintaining key information. This method has been shown to be effective for global homolog detection [16] [8] where the entirety of two proteins correspond to one another. However, it performed worse for the cases where two proteins share all of their domains, but with extended undefined regions between the domains [8]. For local homology detection, where only certain regions of two proteins are similar, this method won’t work well due to the course-grained nature of the representation (as we show in our Results). To remedy these issues we developed a new method (ESM2-RecCut) for predicting domains using the predicted contact maps from ESM-2, and predicted domains are then used for deriving domain-level fingerprints based on iDCT vector quantization.

Protein domains are subunits that can fold and function independently [18]. The protein universe contains many multi-domain proteins, involving a great diversity of domain architectures [19]. A few methods have been developed for protein domain segmentation given protein sequences, including a more recently developed method FUpred (for Folding Unit predictor) [20], which detects domain boundaries from protein sequences based on predicted contact maps. A protein contact map depicts the distances between all residue pairs in a protein, utilizing a binary two-dimensional identity matrix that signifies which pairs are in contact. Sequentially distant residues can be in contact in the tertiary structure. FUpred aims to find domains that maximizes the number of intra-domain contacts, while minimizing the number of inter-domain contacts. For contact map prediction, FUpred generates a multiple sequence alignment (MSA) using the DeepMSA program [21], and the generated MSA is used as the input for contact map prediction using ResPRE [22], a method that couples evolutionary precision matrices with deep residual neural networks. FUpred was shown to outperform existing approaches for contact map prediction, including ConDo [23] and DNN-dom [24]. We found that ResPRE-FUpred pipeline is computationally demanding for creating domain-level DCT fingerprints, and therefore we proposed a new method ESM2-RecCut for this purpose. Our ESM2-RecCut uses contact maps predicted from ESM-2 (together with contextual embeddings of individual residues), taking advantage of its capability of generating contextual embeddings without using MSA so there are no time-consuming iterative searches of similar sequences. In addition, we developed a quadratic time RecCut-based algorithm for domain segmentation given contact map predictions. Predicted domains can then be used for generating domain-level fingerprints to facilitate fast and accurate similarity detection.

## Methods

### Overview of the methods

Our approach, DCTdomain, uses the protein embedding and contact map predictions from ESM-2, detects domains, and represents each protein as one or more DCT fingerprints, including a DCT fingerprint for the whole protein, and if applies, a fingerprint for each of the predicted domains in the protein. The DCT fingerprints are then used for computing the similarity between the proteins. Our method reports two similarity scores: one based on the DCT fingerprints of the whole proteins (denoted as DCTglobal), and the other one based on all DCT fingerprints of whole proteins and of individual domains (denoted as DCTdomain). Figure 1 shows the overview of our approach.

**Figure 1:**
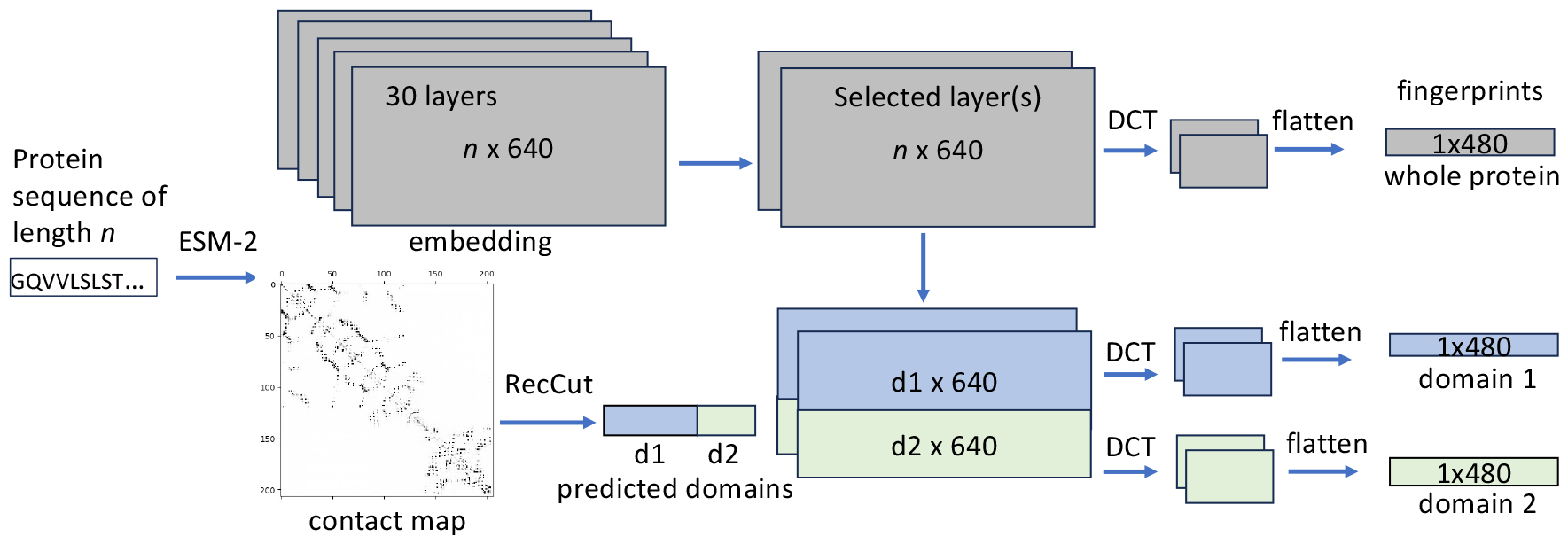
A diagram showing the inference of domain-based embeddings (DCT fingerprints). For the example protein with two domains, three DCT fingerprints will be derived, one representing the whole protein, and the other two are the representations of the domains. This diagram uses a two-domain protein, the ESM-2 t30 model, and fingerprints of size 480 for demonstration purpose without loss of generality.

### Protein language model embedding

We used the output from two layers of the ESM2-t30-150M model (layer 15 and layer 21). It has been shown that each layer of an encoder is able to capture different contextual information about a protein sequence [8]. To find the most informative layers for our particular task, we performed an analysis very similar to HHblits [25], which took pairs of protein domains from SCOPe (v. 2.08) and labeled them as homologous if they belonged to the same fold, and non-homologous if they belonged to different folds. We took the embeddings from every layer of each ESM-2 checkpoint (except for t48, which was too large for our available memory), quantized them, and computed the L1 distance between each protein pair. With these distance values, we calculated the accuracy of homology detection for every layer of each checkpoint. The results were used for selecting the parameters for DCTdomain.

### Protein domain segmentation based on ESM-2 contact map prediction

We used both FUpred [20] and a new recursive cut algorithm (RecCut) to predict domains given predicted contact maps and compared their performance. To distinguish the two versions, we refer to them as ESM2-FUpred and ESM2-RecCut. Comparing to the original FUpred pipeline (which uses ResPRE to predict contact maps, so we refer to it as ResPRE-FUpred), here we used ESM-2’s contact map prediction as input to FUpred or RecCut for domain segmentations. More specifically, ESM-2’s contact map prediction of a protein depicts the probability of any pair of residues i and j being in contact in the protein’s tertiary structure. A discrete version of the contact map referred as *C*[*i, j*] (*C*[*i, j*] = 1 if residue *i* and *j* are in contact; 0 otherwise) is prepared by keeping the top *an* pairs with the highest probabilities (here *n* is the length of the protein, and *a* is a constant, which was empirically tuned using the FUpred benchmark; see below).

We noticed that FUpred/RecCut using ESM-2 contact map prediction results in high precision of the prediction of single domain proteins and high recall for prediction of multi-domain proteins, but tends to split domains into smaller units (see Results). So in some cases, proteins may be segmented into domains or *subdomains*. Domains are typically autonomous structure, function, evolution and folding units of proteins. Domains and smaller subdomains may help reduce the complexity of conformational search by replacing the concerted folding of the entire protein with assembly of folded smaller units [26]. However, we note that the subdomains from ESM2-FUpred or ESM2-RecCut could represent biologically meaningful units (autonomous folding units), or they could result from imperfect contact map prediction where some contacts between the subdomains are not predicted. For simplicity, we don’t distinguish domains and subdomains, and call them domains throughout this paper.

### Recursive cut algorithm for domain segmentation based on contact map

We present the computational problem to partition a given single-chain protein into multiple contiguous or non-contiguous domains, given its contact map (predicted) as the input. Notably, in some cases, the N-terminal and C-terminal of a protein chain are proximal in 3D space, and thus the residues in both terminus can form a single protein domain (e.g., protein 1agr chain A has two domains, one domain contains residues 57 to 177, and the other domain contains two discontinuous regions, 1-56 and 178-350). To account for such cases, herein, we represent the protein as a circular string.

Formally, given a circular string *S*[1, …, *n*] of length *n*, we define a *segmentation* of the string as a sequence of *k* indices (*c*_1_, *c*_2_, …, *c*_*k*_), where 1 *≤ c*_*i*_ *≤ n* represents the indices dividing the string into a set of *k* segments *S*[*c*_*i*_ + 1, …, *c*_*i*+1_] for *i* = 1, 2, …, *k*, and *c*_*k*+1_ = *c*_1_ + *n* to handle the circular nature of the string. We further define a *domain segmentation* of the string as an annotation of each segment by one of its *d* domains, i.e., *l*[*i*] ∈{1, 2, …, *d*} so that all residues in a segment are all assigned to the same domain. So, given a contact map *C*[*i, j*] representing if there is a contact between a pair of residues *i* and *j*, our goal is to find a maximum domain segmentation (i.e., with the maximum number of domains) in which the number of contacts between any two domains is smaller than a threshold. We formulate this problem as the following *domain segmentation problem*.

**Problem 1** (Domain Segmentation Problem). ***Input:*** *A circular string S*[1, …, *n*], *a contact matrix C*[*i, j*] *for* ∀1 ≤ *i, j* ≤ *n, and a threshold W*_*m*_. ***Output:*** *The maximum domain segmentation of S with d domains*, (*c*_1_, *c*_2_, …, *c*_*k*_) *(c*_*k*+1_ = *c*_1_ + *n) and l*[*i*] *∈ {*1, 2, …, *d}, for* 0 ≤ *i* ≤ *k, in which the contact between residues in any two domains is below W*_*m*_, *i*.*e*., *for any i* ≠ *j*, 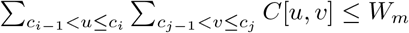

FUpred [20] was previously developed to tackle the computational problem: in each step, an input circular string is partitioned into two contiguous domains (substrings) with the minimum contact, and the procedure continues recursively (i.e., each of the two partitioned substrings forms a circular string that is subject to further partition) until the minimum contact between the two partitioned domains becomes less than the threshold. It was shown that this heuristic approach can reconstruct the known protein domains in most proteins in the curated protein structure classification database SCOP [27], with the exception of rare cases of nested discontinuous multi-domain proteins [20].

Despite the effectiveness of FUpred, it takes *O*(*dn*^3^) running time (where *d* is the number of domains in the protein), which is slow when executed on a large set of proteins, especially those relatively long proteins (e.g., for *n >* 1000). Here, we present the improved algorithm called RecCut (for recursive cut), which follows the heuristic approach to give the same desirable output as FUpred, but running in only *O*(*dn*^2^) time.

Given an input string *S*[1, …, *n*], RecCut considers the 1-cut and the 2-cuts partitions of the protein, where the 1-cut partition at position *i* results in two contiguous, one-segment domains, *S*[1, .., *i*] and *S*[*i* + 1, …, *n*], respectively, while the 2-cuts partition at positions *i* and *j* results in one contiguous, one-segment domain, *S*[*i* + 1, …, *j*], and one non-contiguous domain consisting of two segments, *S*[1, …, *i*] and *S*[*j* + 1, …, *n*], respectively. RecCut aims to first find the 1-cut and 2-cuts partitions of the protein with the minimum contact between two partitioned domains, respectively, and then selects one of them based on two given thresholds (denoted as *W*_1−*cut*_ and *W*_2−*cuts*_, instead of a single threshold *W*_*m*_) to proceed the recursion. Specifically, assuming the between-domain contacts of the 1-cut and 2-cuts partitions are *V*_1−*cut*_ and *V*_2−*cuts*_, respectively, if *V*_1−*cut*_ > *W*_1−*cut*_ and *V*_2−*cuts*_ > *W*_2−*cuts*_, the recursion is terminated and the entire segment is output as a single domain; otherwise, RecCut selects the 1-cut partition (if *V*_1−*cut*_ − *W*_1−*cut*_ ≤ *V*_2−*cuts*_ −*W*_2−*cuts*_) or the 2-cuts partition (if *V*_1−*cut*_ − *W*_1−*cut*_ > *V*_2−*cuts*_ −*W*_2−*cuts*_) for recursion.

The 1-cut protein partition can be achieved in *O*(*n*^2^) time, although a naive approach could take *O*(*n*^3^) time, as for each putative cut site, one needs to compute the contact between the resulting two domains. For the 2-cuts protein partition, a naive approach takes *O*(*n*^4^) time (i.e., for every two putative cut sites, one computes the contact between the resulting two domains). FUpred exploited a dynamic programming (DP) algorithm to reduce it to *O*(*n*^3^). In RecCut, we further optimized the DP algorithm to reduce the running time to *O*(*n*^2^) by introducing three matrices: *E*[*i, j*] representing the sum of the contacts between the residue *i* and all residues within the segment *S*[1, … *j*], *F* [*i, j*] representing the sum of the contacts between the residue *i* and all residues within the segment *S*[*i* + 1, … *j*] and *T* [*i, j*] representing the sum of the contacts between the residues within the segment *S*[*i*, … *j*]. Computation of *E*[*i, j*], *F* [*i, j*] and *T* [*i, j*] for all *i, j ∈*[1, …, *n*] takes *O*(*n*^2^) time, and once computed they can be used to compute *V* [*i, j*], the sum of contacts between the domains resulted from a 2-cuts at *i* and *j*, needed for selecting the best 2-cuts. With the optimization of 1-cut and 2-cuts partition algorithms, RecCut reduces the running time from *O*(*dn*^3^) by FUpred to *O*(*dn*^2^) to compute the domain segmentation, where *d* is the number of domains (typically a small number) and *n* is the protein length. See Figure S1 and details in Supplementary Information for explanation of the RecCut algorithm and the time complexity analysis.

### Compression of protein embedding matrices using iDCT quantization

The iDCT vector quantization method, inspired by DCT-based compression methods, was developed to homogenize the length of vectorized protein sequences while also retaining its sequential pattern and as much information from the original data as possible [16]. It is also a relatively quick computation to perform as the DCT and its inverse are both fast Fourier transforms. This method can reduce both dimensions of an input embedding matrix: 2d-DCT is first applied to the input embedding matrix to compute frequency coefficients, and then an inverse DCT (iDCT) is applied to only low frequency coefficients (discarding the high frequency ones) to produce a dimension reduced matrix, thus achieving compression of the embedding matrix. For example, given an embedded protein sequence from ESM2-t30-150M that has dimensions 250 × 640 (250 residues with each residue embedded as a vector of 640 dimensions), we can reduce both dimensions (e.g., to 3 and 80) that give a compact yet representative representation of proteins for both efficient and accurate homology detection. The compressed matrix is then flattened to produce 1D vector (referred as a *DCT fingerprint*), which represents a protein allowing for the usage of simple vector operations. This way, proteins of various lengths are represented as vectors of the same size, and the vectors are compressed as compared to the original residual level embeddings of the proteins.

For multi-domain proteins, iDCT quantization is applied to each domain to generate a DCT fingerprint for the domain. By doing this, each protein is represented as a DCT fingerprint for single-domain proteins, or as (*d* + 1) DCT fingerprints for proteins with *d* domains. The scipy.fftpack library is used for iDCT quantization, and after iDCT quantization, each number in the vector is multiplied with 127 and saved as an 8-bit integer.

### Protein similarity measure based on DCT fingerprints

Given two DCT fingerprints *DCT*_*i*_ and *DCT*_*j*_ (each of 480 dimensions), their similarity score is defined as *S*(*DCT*_*i*_, *DCT*_*j*_) = 1 −*L*1(*DCT*_*i*_, *DCT*_*j*_)*/c*, where *L*1 is the L1-distance between two vectors and *c* is a normalization constant such that the similarity score is transformed to the range of [0, 1]. By applying the transformation, it is more straightforward to interpret the fingerprint similarity scores and it may be possible to compare similarity scores when fingerprints of various sizes are used. Otherwise, if L1 distances are directly used, they can be of arbitrary scales depending on how iDCT quantization is applied, and the sizes of the fingerprints. In our case for fingerprints of size 480, *c* = 17000. Accordingly, given two proteins *i* and *j* each represented as one (for single-domain proteins) or multiple DCT fingerprints (for multi-domain proteins), their global similarity is computed as the similarity of the DCT fingerprints of the whole proteins, and their local similarity is computed as the maximum similarity of any pairs of DCTs (including those for the whole protein and those for individual domains) from the two proteins. We note that L1-distance was used in PROST to quantify the distance between DCT matrices. Our similarity score is based on L1-distance, but it is transformed and normalized to be in the range of [0, 1] (0 for no similarity).

## Benchmarks

We used the FUpred benchmarks [20] for testing domain segmentation using ESM-2 contact maps. Similarly, we used the train set (with 2549 proteins) to tune the parameters including *W*_1−*cut*_ and *W*_2−*cuts*_ (corresponding to Cutoff2c and Cutoff2d in FUpred [20]), for distinguishing between continuous multi- and single-domain proteins, as well as discontinuous multi- and single-domain proteins, respectively, and reported the performance on the test collection with the same number of proteins. The train and test collections don’t share proteins. The benchmark was downloaded from https://zhanggroup.org/FUpred/.

For homolog detection, we first used a curated benchmark of protein pairs [28] which labels proteins as homologous if their domains, in consecutive order, are in the same family/clan/superfamily, and non-homologous if no domain in the first protein is a part of the same family/clan/superfamily as any domain in the second. All proteins were derived from a given specific set of genomes (16 species including three prokaryotes and 13 eukaryotes). Like in PROST [8], we used two groups of this benchmark: max50, where the maximum distance between two domains is 50 residues, and nomax50, where there is no limit between domains. These benchmarks are further divided into subsets based on which database the domains came from - pfam[29], CATH[30], and SCOP[31]. PROST [8] showed that the max50 datasets were easier to perform well on than the nomax50 datasets, so we focus our analysis mostly on the nomax50 groups. See Table 1 for a summary of each benchmark and example cases of homologs in each group. We re-tested each homolog detection method (including HHsearch [6], UBLAST [32], FASTA [33], phmmer [5], BLAST [2], and CS-BLAST [4]) on each dataset and found nearly identical Area Under Curve (AUC) values as originally reported [28]. We note that we built profiles using HHsearch from scop40 to achieve better sensitivity than what was originally reported. This also caused an increase in search time. The benchmark was downloaded from http://sonnhammer.org/download/Homology_benchmark.

**Table 1:** Protein benchmarks for homology/similarity detection.

| Benchmark * | # pairs<br>(homologs) | homolog definition | example protein [domain architecture] |
| --- | --- | --- | --- |
| pfam-max50 | 10450<br>(5228) | identical domain architecture;<br>< 50 aa between domains | Q9VFJ2 [PF03946,PF00298]<br>P53875 [PF03946,PF00298] |
| pfam-nomax50 | 71988<br>(36278) | identical domain architecture;<br>no constraint on the aa between<br>domains | Q15149 [PF03501,CL0188,CL0188,PF00681]<br>Q9QXS1 [PF03501,CL0188,CL0188,PF00681] |
| pfam-local | 15273<br>(7602) | share some domains, but not all | P40791 [PF00319,**PF12347**]<br>Q8VWM8 [PF00319,**PF01486**] |
| gene3d-nomax50 | 58163<br>(29109) | same as pfam-nomax50<br>but based on CATH domains | P52917 [1.20.58.280,3.40.50.300]<br>Q9ZNT0 [1.20.58.280,3.40.50.300] |
| supfam-nomax50 | 49365<br>(24708) | same as pfam-nomax50<br>but based on SCOP domains | Q9T0N8 [56176,55103]<br>P46681 [56176,55103] |

\* The benchmarks are denoted as pfam-max50, gene3d-nomax50 and so on to indicate the domain database used for defining the homologs. The benchmarks include full length proteins.

Based on the pfam-nomax50 benchmark, we further created a benchmark for testing detection of local similarities between proteins (refered as pfam-local). This benchmark contains homologs that share at least some of their domains, but not all. An example is shown in Table 1: the two proteins each have two domains and they share one domain (PF00319).

For performance evaluation of domain segmentation on the FUpred benchmark, we focused on the accuracy of classifying proteins into single-domain versus multi-domain proteins, and used various metrics for evaluation, including accuracy and Matthew’s correlation coefficient. For evaluation of homology detection, we calculated the AUC from the ROC plots to show the true positive rates vs false positive rates for the different methods. Each of the benchmarks contains a nearly equal number of homologs (positives) and non-homologs (negatives); see Table 1.

## Availability of data and code

Our programs (RecCut for domain domain segmentation in C++, and Python scripts for sequence embedding, DCT fingerprint generation, and sequence comparison and similarity search based on DCT fingerprints) are available as open source on github at https://github.com/mgtools/DCTdomain. Benchmarks, results, and the command-line calls for each homology detection tool are also available in the same github repo.

## Results

### Domain predictions using predicted contact maps from ESM-2

Comparing to FUpred predicted domains using predicted contact maps from ResPRE (ResPRE-FUpred), our approaches that use contact maps from ESM-2 (ESM2-FUpred, and ESM2-RecCut) gave very good recall of the predictions for multi-domain proteins and precision for single-domain proteins, as shown in Table 2, however, their overall accuracy was lower. The results also suggested that approaches based on ESM-2 contact maps tend to split domains into smaller units (some single-domain proteins were predicted to be multi-domain proteins so more domains are predicted for multiple-domain proteins). Proteins in the test dataset of 2549 proteins each contain 2.47 domains on average, and ESM2-FUpred and ESM2-RecCut predicted 3.21 and 3.03 domains per protein on average, respectively. Figure 2 shows two examples of ESM2-RecCut predictions. In the first case, 1a04a, ESM2-RecCut gave almost identical domain predictions with 3D structure based domain segmentation in SCOP (with the boundary of the domains shifted by one residue). Visualization of the contact map from ESM-2 shows that this protein has two domains. By contrast, ESM2-RecCut predicted two domains in d1wd3a1, although SCOP considers it a single-domain protein. We also note that among the 2549 proteins in the benchmark, 133 proteins contain discontinuous domains. ESM2-RecCut predicted 67 of these proteins having discontinuous domains. It is likely that the contacts between residues from the segments in the discontinuous domains are more difficult to predict, and therefore discontinuous domains could be underestimated by ESM2-RecCut.

**Table 2:** Single- and multi-domain classification results on 2549 test proteins from the FUpred benchmark.

| Method | Multi-domain |  | Single-domain |  | All |  |
| --- | --- | --- | --- | --- | --- | --- |
|  | Precision | Recall | Precision | Recall | ACC | MCC |
| ResPRE-FUpred* | 0.860 | 0.873 | 0.936 | 0.929 | 0.910 | 0.799 |
| ESM2-FUpred | 0.631 | 0.974 | 0.982 | 0.716 | 0.802 | 0.651 |
| ESM2-RecCut | 0.663 | 0.941 | 0.963 | 0.761 | 0.821 | 0.663 |

\*ResPRE-FUpred results are taken from [20]. In principle, ESM2-FUpred and ESM2-RecCut should result in the same results as they use the same scoring scheme (so called FUscore). However, there are certain technical details that we cannot replicate in ESM2-RecCut so the results vary slightly. ‘ACC’ and ‘MCC’ are the accuracy and Matthew’s correlation coefficient, respectively. The results shown here were based on contact map predictions using the ESM-2 t30 model. Refer to Supplementary Table S1 for the results based on contact map predictions using the ESM-2 t33 model, which gave more accurate domain predictions but performed worse for our task of homology detection.

**Figure 2:**
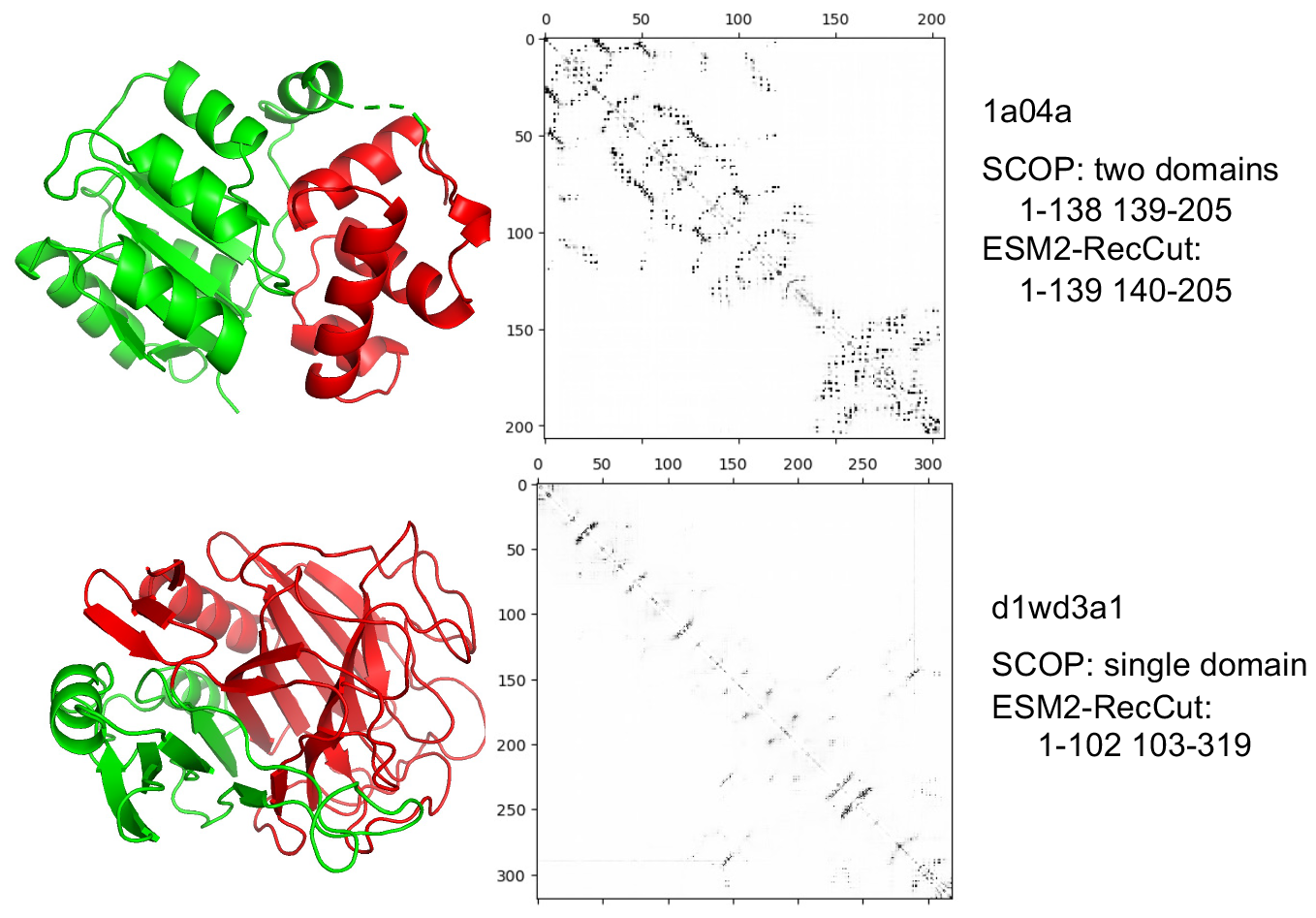
Examples of domain segmentation using ESM2-RecCut. The cartoon visualizations (by PyMol) of the protein structures are shown on the left, with predicted domains (by ESM2-RecCut) shown in different colors: the first domain in green and the second domain in red. The figures in the middle show ESM-2 contact maps of the two proteins.

We were unable to directly compare the running time of the different methods, as we couldn’t run ResPRE to predict contact maps (the FUpred package lacks some of the important files needed for running the pipeline). It was reported in the paper [20] that it takes a few hours on average to predict domains for a protein which is longer than 400 amino acids. The ResPRE-FUpred method is time consuming because it involves two slow steps: the contact map prediction step (ResPRE) and the domain prediction (FUpred) in cubic time. Our method ESM2-RecCut reduces the running time significantly by utilizing ESM-2’s contact map prediction (it was shown that by bypassing the generation of MSA, ESMFold achieved an order-of-magnitude acceleration comparing to AlphaFold2 [13]). In addition, RecCut reduces the time complexity for domain segmentation from cubic time to quadratic, with respect to the protein length.

Here we showed the comparison of running time spent on the domain segmentation step (not including the contact map prediction) by the different approaches. For the 2549 FUpred benchmark proteins, FUpred ran in 158 seconds, whereas RecCut took 40 seconds. The running time difference is more significant on the homolog detection benchmarks (with much longer proteins than the FUpred benchmark). For example, for a total of 13342 proteins from the pfam-nomax50 benchmark (some of the proteins are very long; for example, the longest protein Q91ZU6 has 7393 residues), given their predicted contact maps (by ESM-2), it took FUpred 32 hours to predict the domains, whereas it took RecCut only 18 mins. Supplementary Figure S2 plots the running time of RecCut versus protein length, showing a quadratic relationship consistent with the theoretical analysis. This figure also shows the number of domains versus protein lengths, suggesting a linear relationship with roughly one domain per 100 residues.

Combining the accuracy and running time, we chose to use ESM2-RecCut as the approach for predicting domains, and used it for preparing domain level DCT fingerprints for homology detection. We show below that ESM-2 contact map based domain predictions, although not as accurate as those from ResPRE-FUpred, still gave good performance for using DCT fingerprints to detect similarity between proteins.

### Selection of parameters for DCT fingerprint construction

We tested the impact of using different ESM-2 layers from different models (checkpoints) on the performance of homology detection based on DCT fingerprints. For this testing, we used domains defined in SCOPe (see Methods) as the inputs so no domain segmentation was involved (which otherwise would have impacts on the performance of our approaches as well). Figure 3 summarizes the results, showing that using ESM-2 t30 model (with 150M parameters) and t33 model (with 650 parameters) resulted in better homology detection than using the other models, and using layers 15 and 21 from t30 resulted in the best results than other layers. On the embeddings produced by these layers we tested different dimensions for iDCT quantization and found that reducing each embedding matrix to 3×80, which was the smallest representation of embeddings while retaining the most information in our tests (results found in Supplementary Figure S3). For comparison, PROST uses ESM1b and layers 14 and 26 to derive DCT fingerprints of size 475 [8].

**Figure 3:**
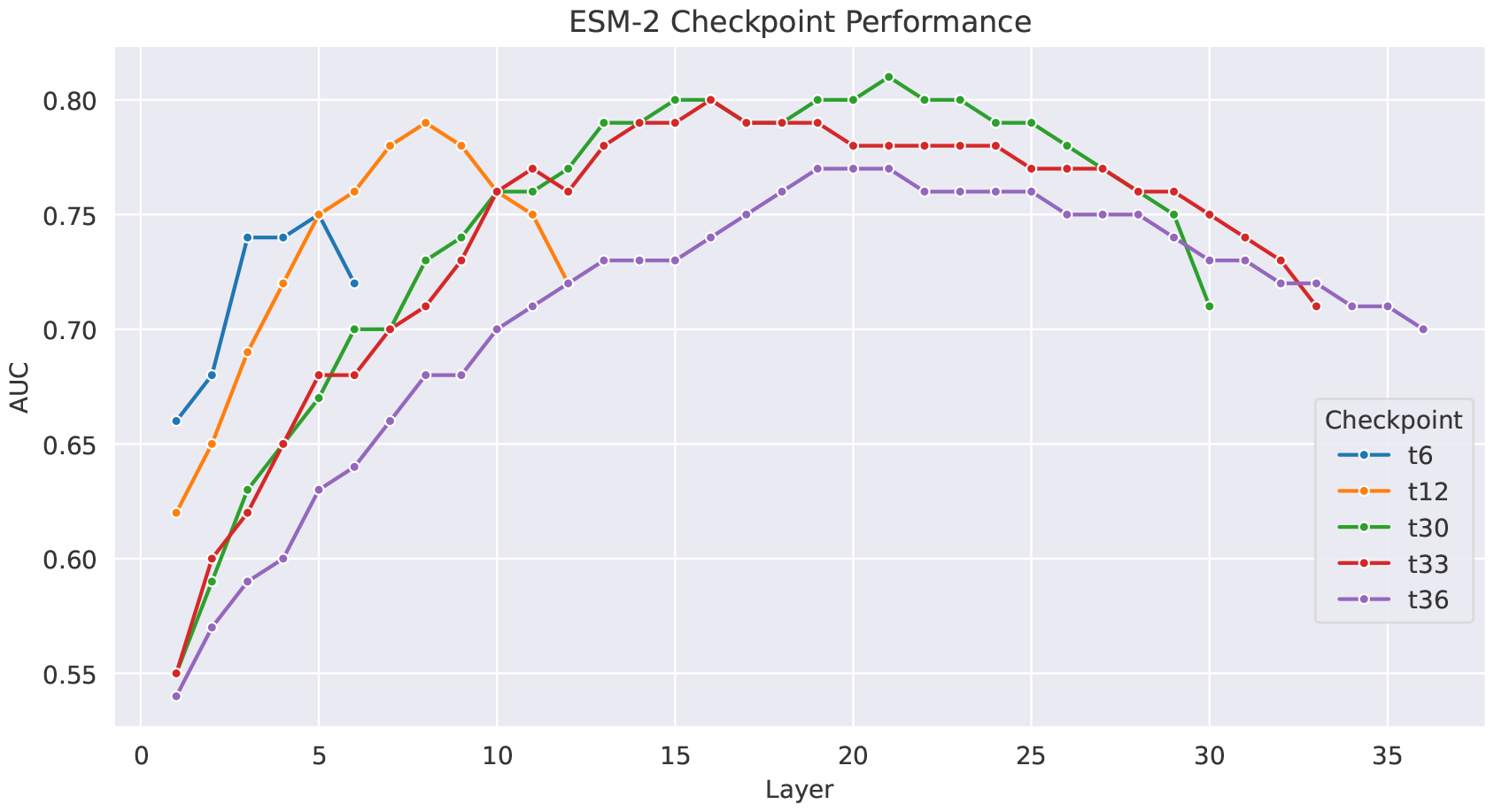
Comparison of the performance of every ESM-2 checkpoint (except t48) by layer on determining if sequence pairs from SCOPe v2.08 are within the same fold or different fold.

Considering that using ESM-2 t30 for embedding proteins resulted in best homology detection, despite that the bigger model t33 resulted in better domain segmentation, we chose to use ESM-2 t30 as the default model for our pipeline. All the DCTdomain and DCTglobal results shown below were based on DCT fingerprints from two layers of the ESM-2 t30 embeddings of proteins: layers 15 and 21 each transformed into a 3×80 matrix, and the two reduced matrices are flattened and concatenated into a vector of 480 dimensions.

### Results on global homolog benchmarks

We applied our method, DCTdomain, which computes the similarity between proteins using their DCT fingerprints (of individual domains and whole proteins) to the four global homolog benchmarks, and compared the results with using the DCT fingerprint of the whole proteins only (DCTglobal and PROST) and other methods for homology detection. We note DCTglobal is essentially the same as the PROST method from algorithmic perspective (both using DCT fingerprints of whole proteins); however, their performance varied slightly due to the small differences of their implementation details (ESM model and layers used, and the size of the DCT fingerprints). For calculating the AUC, we used the DCT similarity score (see Methods) for DCTdomain and DCTglobal, the L1-distance between DCT fingerprints for PROST, and bit-scores for all the other sequence/profile based methods that we compared.

Figure 4 shows the comparison of homology detection on the four benchmarks (pfam-max50, pfam-nomax50, gene3d-nomax50, and supfam-nomax50). Our results showed that using DCT fingerprints of whole proteins (PROST and DCTglobal) underperformed, achieving worse AUC values than a few methods on the nomax50 benchmarks (although they still achieved a good performance with AUC values higher than FASTA and UBLAST). This result is consistent with [8] that the global DCT based distance didn’t work well for detecting similarity between protein pairs that share global similarities but with extended, undefined regions between shared domains. By contrast, DCT similarity based on domains (DCTdomain) maintained better results than any other method, including DCTglobal and profile methods like HHsearch and CS-BLAST. This is particularly true for sequences from the structural databases, SCOP (supfam) and CATH (gene3d), where DCTdomain outperforms HHsearch in AUC as much as HHsearch outperforms the next best methods (phmmer and CS-BLAST).

**Figure 4:**
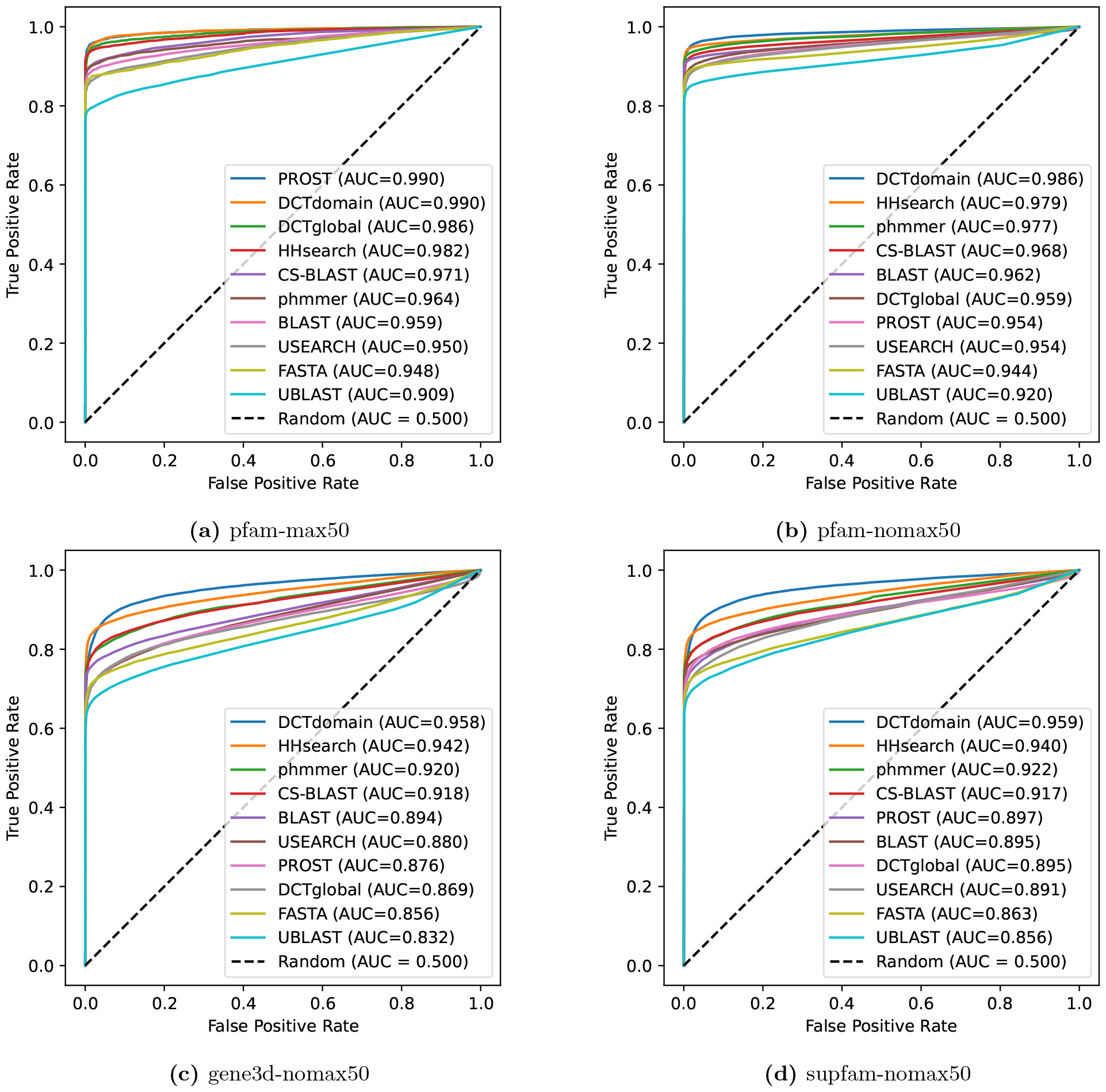
ROC plots for comparison of the different methods on four benchmarks of global homologs. pfam-max50 benchmark; (b) pfam-nomax50 benchmark; (c) gene3d-nomax50 benchmark; (d) supfam-nomax50 benchmark.

### Results on local homolog benchmark

Figure 5 shows that DCTdomain works well for detecting local similarities between the proteins (the proteins don’t share global similarities). DCTdomain performed as well as HHsearch on this benchmark and clearly outperformed all other methods. Given that a DCT fingerprint is an averaged representation, it is not surprising that DCTglobal and PROST had the worst performance on this benchmark.

**Figure 5:**
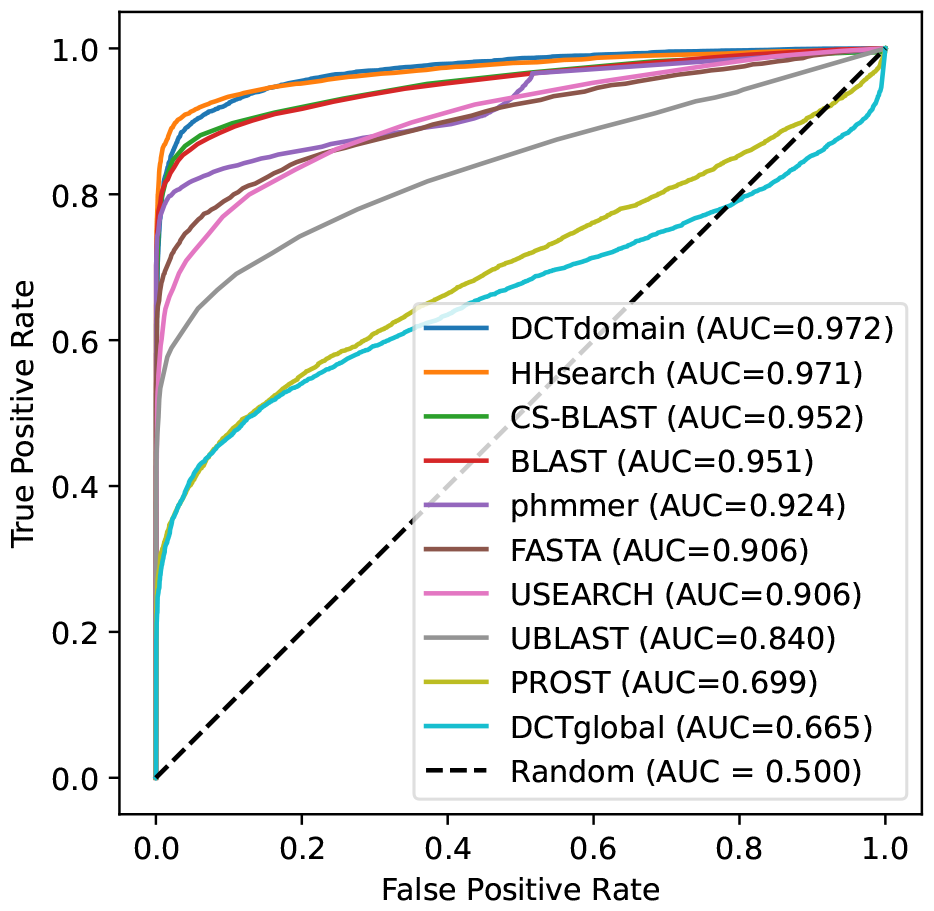
ROC plots for comparison of the different methods on the pfam-local benchmark that contains proteins sharing local similarities.

Table 3 summarizes the running time of the different approaches along with their performance in AUC. The results show that using DCT fingerprints achieved fast similarity calculation comparing to other methods. Although our approach DCTdomain performed roughly equivalent to HHSearch in AUC on this benchmark, our method is more than three orders of magnitude faster (without including the DCT fingerprint preparation time). DCTdomain is still significantly faster, even when including the time for DCT fingerprint generation. We note the times reported in Table 3 don’t include the time for construction of the search database that is required by some of the other approaches (BLAST, CS-BLAST, HHsearch, and UBLAST), and HHsearch was very slow due to the profile construction step. We acknowledge that ESM-2 embedding and domain segmentation take time especially for long proteins (see Supplementary Figure S4), but in practice we only run the calculation once, and the DCT fingerprints of proteins can be computed and saved in a numpy compressed NPZ file for later applications.

**Table 3:**
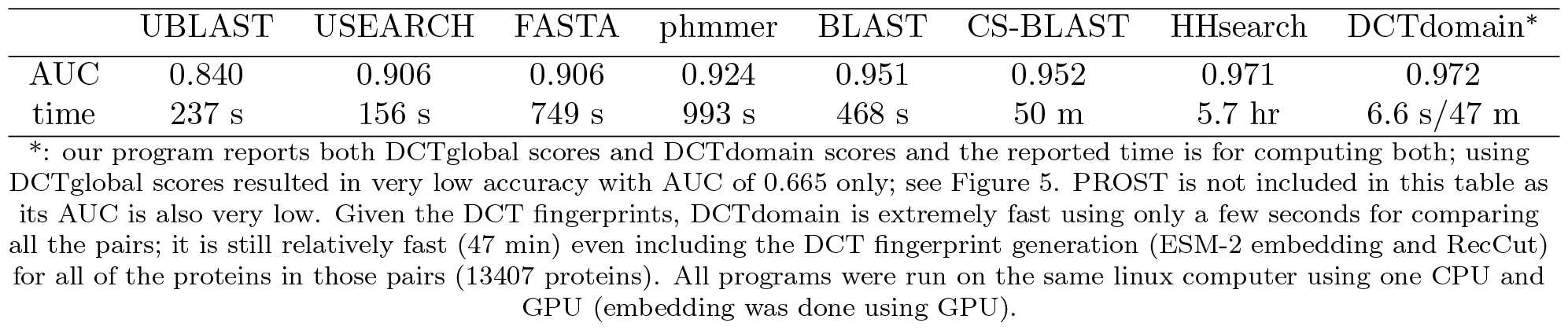
AUC and total runtime for each method (listed from the least to most accurate) on pfam-local benchmark with 15273 pairs of proteins.

### Examining the difference between DCTdomain and DCTglobal using G6PD containing proteins

Results shown above clearly demonstrated the difference of the DCTdomain and DCTglobal scores for detecting similarity between mult-domain proteins. Here, we applied DCTdomain and DCTglobal to a collection of G6PD containing proteins considering that G6PD is found in various domain architectures and G6PD-containing proteins have important functions. G6PD proteins (glucose-6-phosphate dehydrogenase) are enzymes whose main function is to produce NADPH, a key electron donor in the defense against oxidizing agents and in reductive biosynthetic reactions. We used InterPro [34] to look up G6PD N (PF00479, Glucose-6-phosphate 1-dehydrogenase, NAD-binding domain) containing proteins, which showed that this domain can be found in 163 different domain architectures, among which, the domain architecture (G6PD N - G6PD C) is the dominate one found in 44k sequences (the second most frequent domain architecture is found in 1658 proteins). We collected a total of 28 G6PD containing proteins representing different domain architectures, including 8 sequences that have this domain architecture: PF00479 (G6PD N) - PF02781 (G6PD C) PF02781 (G6PD C) - PF01182 (Glucosamine iso), 7 sequences with this domain architecture PF00479 PF02781 - PF13347 (MFS 2), 6 sequences of this architecture PF03446 (NAD binding 2) - PF00393 - PF00479 - PF02781, and 6 sequences of this domain architecture PF00479 - PF02781 - PF08123 (DOT1). Since these proteins all contain G6PD domain, pairs of these proteins can be global homologs (they share the same domain architecture), or share local similarities (some domains are different). We compared the distributions of the DCT fingerprint similarity of the global homologs, partially similar pairs, and non-homologs (from the pfam-nomax50 collection). Figure 6 shows that all similar proteins (sharing global or local similarities) have high DCTdomain similarity, forming distinct distributions that are well separated from non-homolog DCT similarity distribution (Figure 6a). By contrast, if using DCTglobal similarities, the separation between the DCT similarity distributions of the local homologs and the non-homologs diminished (Figure 6b).

**Figure 6:**
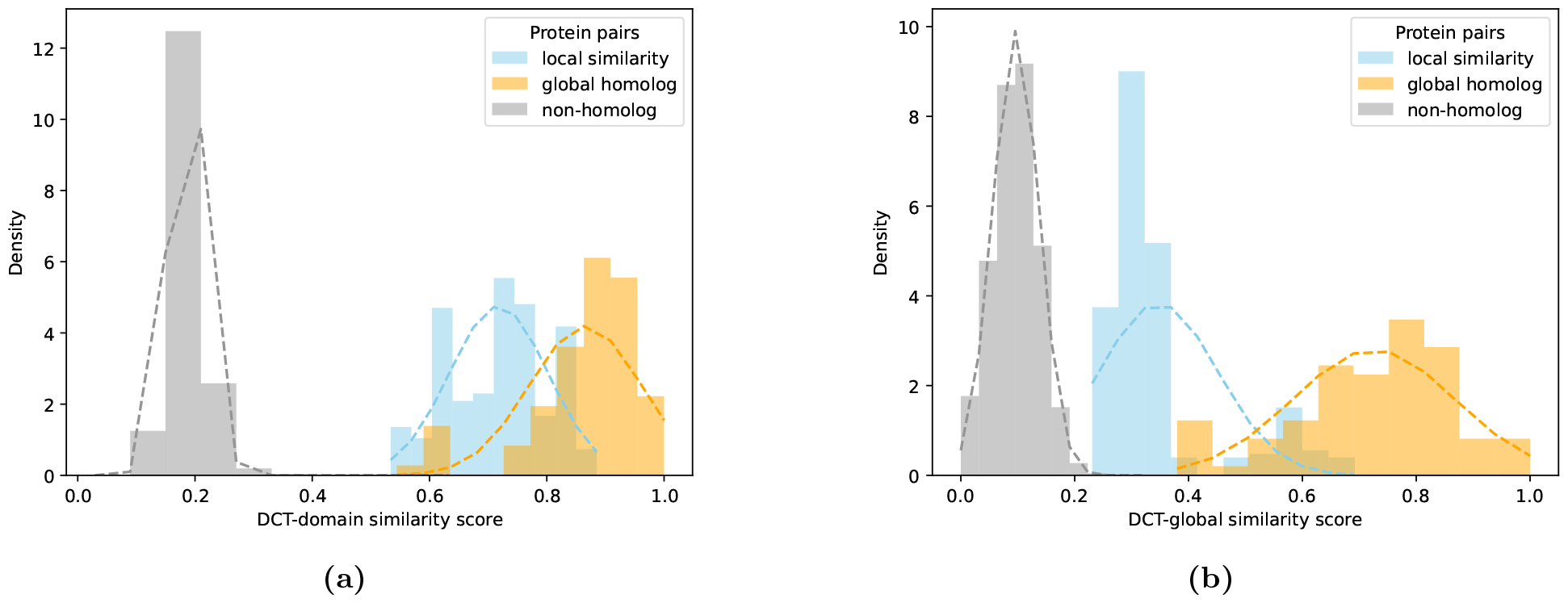
Comparison of domain embedding based similarity between G6PD containing proteins. (a) DCTdomain; (b) DCTglobal.

## Discussion

Using ESM-2 to predict contact maps for subsequent domain prediction with RecCut, as well as generating embeddings that were compressed using iDCT quantization, proved to be an effective method for detecting sequences that are homologous. DCTdomain performed well on every benchmark we tested it against. It performed the best on both global homolog benchmarks relative to every other method we tested, particularly for structure-based classifications, and was as sensitive as HHsearch on the local homolog benchmark. Using the FUpred domain prediction benchmark, it is clear that ResPRE-predicted contact maps are more effective than ESM2-predicted contact maps for domain prediction. However, our method ESM2-RecCut is significantly faster than the existing approaches that rely on more accurate but slower contact map prediction methods. In addition, as our results showed, the domain segmentations achieved by ESM2-RecCut, despite imperfect, are already very helpful for generating DCT fingerprints for local homology detection. We anticipate that our RecCut approach can also be used for automatic domain segmentation when 3D structures of proteins are available. We note there are recent developments of algorithms for domain parsing given tertiary structures of proteins (real or model) including DPAM [35] and Unidoc [36]. We will look into the possibility of adapting some of these methods into our DCTdomain pipeline.

Since multi-domain proteins are prevalent, and various domain architectures found in those proteins have important structural and functional implications, it is important to develop methods that can effectively compare multi-domain proteins. We showed that it is important to have domain or sub-domain level representations such as the DCT fingerprints for homology detection, and we proposed a method that utilizes domain segmentation based on contact maps for this purpose. In this work, we compute the similarity of two proteins as the highest similarity of any two DCT fingerprints of the domains found in the proteins, and we anticipate that other metrics may be developed for more sensible similarity quantification.

Our tests showed that using the ESM-2 t33 checkpoint for contact map predictions resulted in better domain segmentation than using t30 (see Table 2 and Table S1 for a comparison). However, using domain predictions based on contact maps from the larger t33 model resulted in slightly worse homology detection. Since homology detection is the main goal of this paper, in combination with that t30 is a much simpler model with 150M parameters comparing to t33 with 650M parameters so it is more memory efficient, we chose t30 as the default checkpoint for DCTdomain. However, if a user is interested in using ESM2-RecCut for domain prediction purpose, we would recommend using t33, which is also available as an option in our pipeline. There are also other parameters involved for the domain segmentation based on ESM-2 contact maps including the thresholds *W*_1−*cut*_ and *W*_2−*cuts*_. If the users want to apply our RecCut algorithm but use different language models, they may have to re-tune those parameters using a benchmark such as the FUpred benchmark as we did.

We note ESM-2 embedding of long proteins is memory extensive. We used a simple strategy to dissect long proteins into overlapping segments, embed individual segments, and then use the average of the embeddings and contact maps for the overlapping regions. Our tests showed that using such a strategy resulted in very similar results for homology detection with must faster embedding times since embedding is normally non-linear in relation to sequence length.

We anticipate that DCT fingerprints can be applied to other applications. They can be used to enable fast database searches, where DCT fingerprint based searches can be combined with sequence-based methods (such as MMSeq2 as in knnProtT5 [7]) or alignment methods that utilize contextual embeddings of individual residues (PEbA [37] or vcMSA [38]) for more accurate and sensitive detection of similar proteins. They may also be applied for other structural and functional analysis of proteins.

