## Supplementary Information for "Protein domain embeddings for fast and accurate similarity search"

#### RecCut algorithm

Given a protein (represented as a string $S[1, ..., n]$) and its predicted contact map $C[i, j]$ for $i, j \in [1, \ldots, n]$ ($C[i, j]$ is 1 if there is a contact between residue $i$ and $j$, 0 otherwise), the protein segmentation is to partition it into domains such that contacts within domains are maximized and contacts between domains are minimized. The problem can be solved using a recursive approach, called RecCut (see Algorithm 1 below). In each iteration, RecCut checks 1-cut partition (Algorithm 2), and 2-cuts partition (Algorithm 3), and decides if the segment shall be reported as a single domain (so the recursion terminates), or two domains resulted from a 1-cut, or 2-cuts.

Figure S1 gives a schematic demonstration of the algorithms.

The 1-cut protein partition can be solved in $O(n^2$ using the following recursive formula:

$$V[i] = V[i-1] - G[i] + H[i] \tag{1}$$

where $V[i]$ is the sum of contacts between domains from 1-cut at $i$, $H[i]$ is the sum of contacts between residue $i$ and residues in $S[1, \ldots, i]$, and $G[i]$ is the sum of contacts between residue $i$ and residues in $S[i+1, \ldots, n]$.

The 2-cut protein partition can be solved in $O(n^2)$ using the following recursive formula:

$$\begin{aligned} E[i,j] &= E[i,j-1] + C[i,j] \\ F[i,j] &= E[i,j] - E[i,i-1] \\ T[i,j] &= T[i+1,j] + F[i,j] \\ V[i,j] &= T[1,j] + T[i,n] - T[1,i] - T[j,n] - 2 \cdot T[i,j] \end{aligned} \tag{2}$$

where $V[i, j]$ is the sum of the contacts between two domains from 2-cuts at $i$ and $j$, $E[i, j]$ is the sum of the contacts between residue $i$ and all residues within $S[1, \ldots, j]$, $F[i, j]$ is the sum of the contacts between residue $i$ and all residues within $S[i+1, \ldots, j]$, and $T[i, j]$ is the sum of the contacts between the residues within S[i,. . . ,j], and $V[i, j]$ is the sum of the contacts between the residues from the two domains.

**Algorithm 1** The RecCut algorithm for recursive domain segmentation

```
function RecCut(S[1,...,n], n, W_{1-cut}, W_{2-cuts}, C[1...n,1...n])
    V_{1-cut}, i ← OneCutPartition(S[1,...,n], n, C[1...n,1...n])
    V_{2-cuts}, i, j ← TwoCutsPartition(S[1,...,n], n, C[1...n,1...n])
    if V_{1-cut} > W_{1-cut} and V_{2-cuts} > W_{2-cuts} then
        Output S[1,...,n]
    else if V_{1-cut} − W_{1-cut} ≤ V_{2-cuts} − W_{2-cuts} then
        RecCut(S[1,...,i], i, W_{1-cut}, W_{2-cuts}, C[1...n,1...n])
        RecCut(S[i+1,...,n], n − i, W_{1-cut}, W_{2-cuts}, C[1...n,1...n])
    else
        RecCut(join(S[j+1,...,n], S[1,...,i]), n − j + i, W_{1-cut}, W_{2-cuts}, C[1...n,...n])
        // join() concatenates two segments
        RecCut(S[i+1,...,j], j − i, W_{1-cut}, W_{2-cuts}, C[1...n,1...n])
    end if
end function
```

**Algorithm 2** The 1-cut algorithm

```
function OneCutPartition(S[1,...,n], n, C[1...n,1...n])
    G[1] ← 0
    for i ← 2 to n do
        G[i] ← G[i − 1]
        for j ← 1 to i − 1 do
            G[i] ← G[i] + C[j, i]
        end for
    end for
    H[n] ← 0
    for i ← n − 1 downto 1 do
        H[i] ← H[i − 1]
        for j ← i + 1 to i do
            H[i] ← H[i] + C[i, j]
        end for
    end for
    V[1] ← 0
    MinV ← ∞
    for i ← 2 to n do
        V[i] ← V[i − 1] − G[i] + H[i]
        if V[i] < MinV then
            MinV ← V[i]
            MinIndexI ← i
        end if
    end for
    Return MinV, MinIndexI
end function
```

**Algorithm 3** The 2-cuts algorithm

```
function TwoCutsPartition(S[1,...,n], n, C[1...n,1...n])
    for i ← 1 to n do
        E[i,0] ← 0
        for j ← 1 to n do
            E[i,j] ← E[i,j−1] + C[i,j]
        end for
    end for
    for i ← 1 to n do
        for j ← i+1 to n do
            F[i,j] ← E[i,j] − E[i,i−1]
        end for
    end for
    for i ← 1 to n do
        for j ← i+1 to n do
            T[i,j] ← 0
        end for
    end for
    for j ← 1 to n do
        for i ← j−1 downto 1 do
            T[i,j] ← T[i+1,j] + F[i,j]
        end for
    end for
    MinV ← ∞
    for i ← 1 to n do
        for j ← i+1 to n do
            V[i,j] ← T[1,j] + T[i,n] − T[1,i] − T[j,n] − 2 · T[i,j]
            if V[i,j] < MinV then
                MinV ← V[i,j]
                MinIndexI ← i
                MinIndexJ ← j
            end if
        end for
    end for
    Return MinV, MinIndexI, MinIndexJ
end function
```

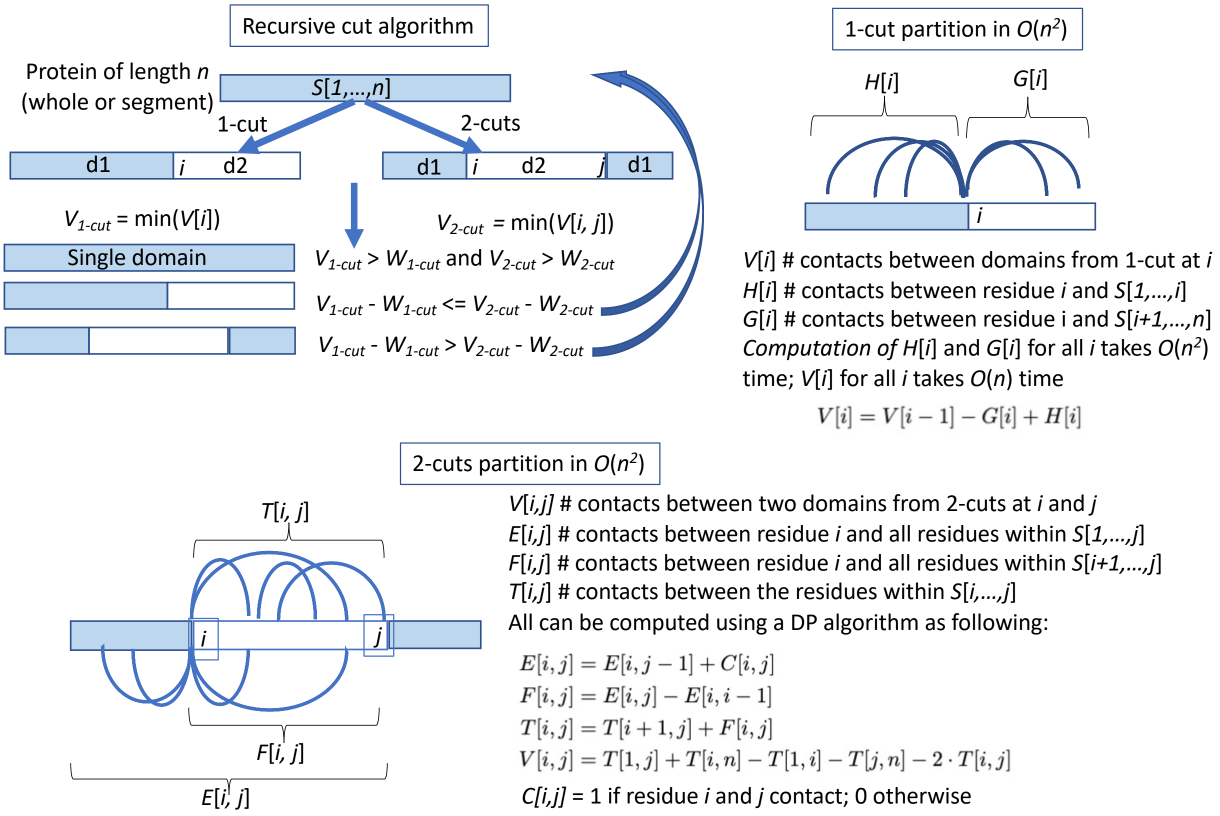


Figure S1: Schematic demonstration of the RecCut algorithm, and algorithms for the 1-cut and 2-cuts protein partition problems. The arcs in the plots represent contacts betweent the residues.

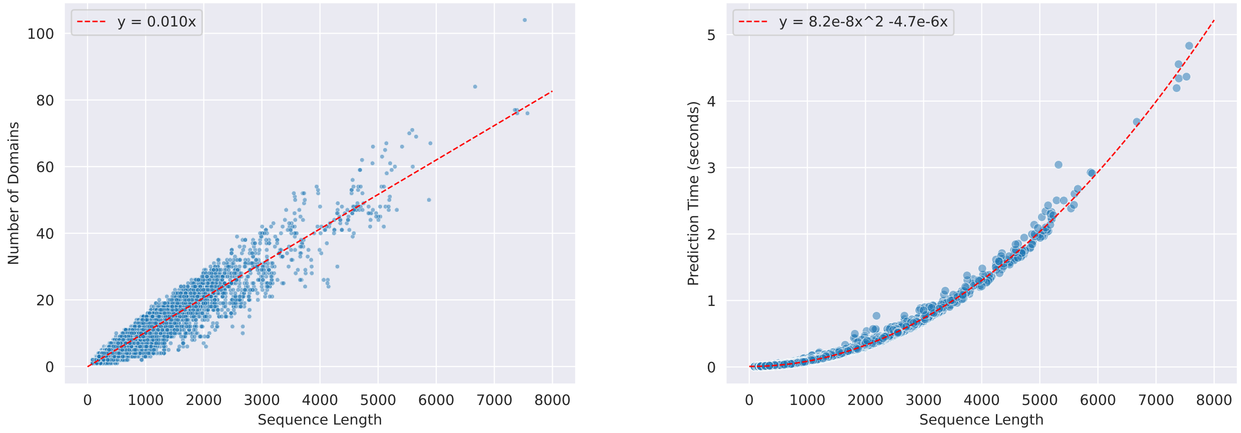


(a) Number of domains per protein

(b) RecCut running time versus protein length

Figure S2: Number of predicted domains and running time for proteins of various lengths.

### RecCut Results

We tested the impacts of using different ESM-2 models on the domain predictions. Table S1 shows the results of RecCut on FUpred on the benchmark when ESM-2 t33 was used (instead of t30). The results showed that using the larger t33 model gave more accurate domain predictions.

Table S1: Single- and multi-domain classification results on 2549 test proteins from the FUpred benchmark.

| Method | Multi-domain | | Single-domain | | All | |
|---|---|---|---|---|---|---|
| | Precision | Recall | Precision | Recall | ACC | MCC |
| ResPRE-FUpred | 0.860 | 0.873 | 0.936 | 0.929 | 0.910 | 0.799 |
| ESM2-FUpred | 0.700 | 0.939 | 0.963 | 0.799 | 0.846 | 0.700 |
| ESM2-RecCut | 0.729 | 0.890 | 0.938 | 0.835 | 0.853 | 0.696 |

'ACC' and 'MCC' are the accuracy and Matthew's correlation coefficient, respectively. The results shown here were based on contact map predictions using ESM-2 t33. Refer to Table 2 in the main paper for the results based on contact map predictions using ESM-2 t30.

We also empirically tested the time complexity of RecCut using proteins of various lengths. We combined all sequences in the three nomax50 datasets (pfam, gene3d, and supfam) for a total of 18876 unique sequences with an average length of about 800 amino acids per sequence. With each sequence's predicted contact map, we ran RecCut to see how sequence length affected the number of domains predicted and the time it took to run the program. Our results show that for a sequence of length 1000, about 10 domains are predicted, with a linear increase in domains as sequence length increases. For a sequence of the same length, it takes RecCut about 0.0004 seconds on an AMD EPYC 74F3 to segment the sequence, with a quadratic increase in time as sequence length increases. We visualize our results and provide the linear and quadratic regression equations for the number of domains predicted and prediction time, respectively, in Figure S2.

### iDCT Quantization Dimensions

After finding that layers 15 and 21 from the ESM2-t30-150M model provided the most informative embeddings for our task of homology detection, we performed the same analysis using protein pairs from SCOPe (v 2.08) to determine which dimensions for iDCT quantization of embeddings were able to preserve the most

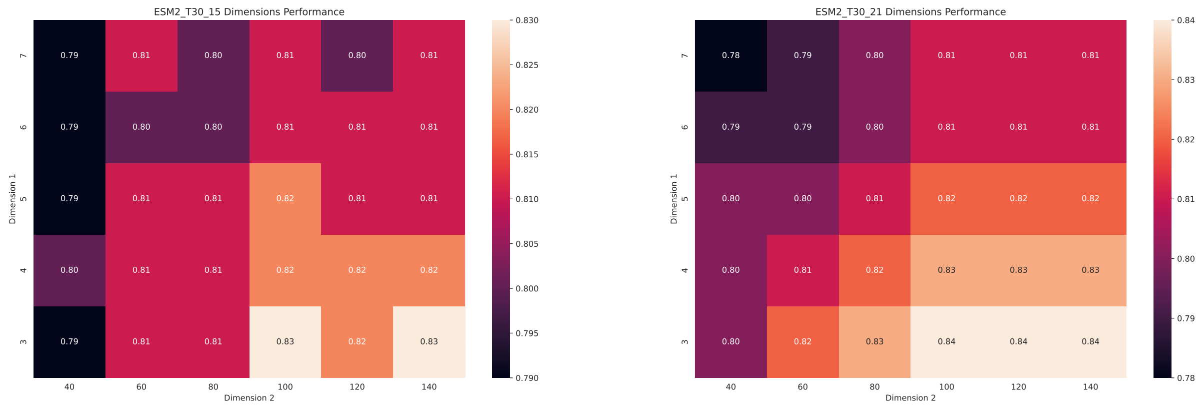


(a) ESM2-t30-layer 15

(b) ESM2-t30-layer 21

Figure S3: Heatmaps showing the impacts of the dimensions of iDCT transformation of the residue embeddings as measured by AUC.

information from these embeddings while keeping fingerprint size relatively small. Figure S3 shows that representing both layers in small matrices of 3x80 gave good results.

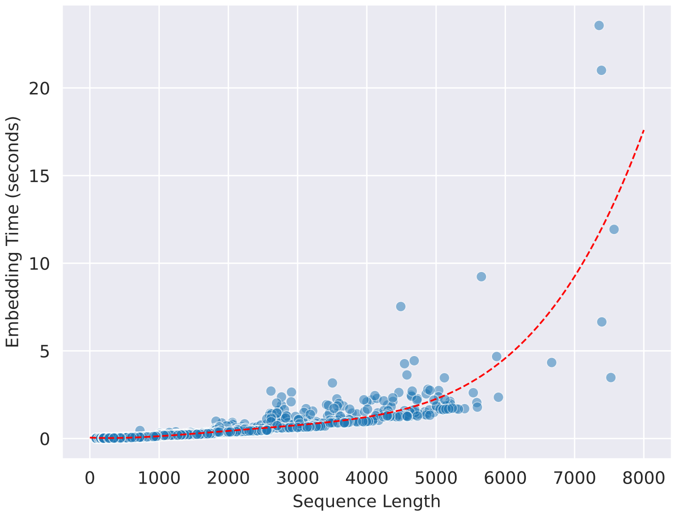


Figure S4: Embedding time using ESM2-t30-150M for sequences from all nomax50 datasets (18876 sequences). Embedding of short sequences is very fast, about 0.05 seconds for proteins of length less than 1000 residues. We noticed a few long protein outliers which took substantial time to embed.
